# Epo and hypoxia accelerate a pattern of gradual cell cycle shortening in BFU-e and CFU-e erythroid progenitors *in vivo*

**DOI:** 10.1101/2025.11.18.688855

**Authors:** Ashley Winward, Logan Lalonde, Divya Nair, Merav Socolovsky

## Abstract

Regulation of the earliest erythroid progenitors is not well understood, yet it is relevant to some types of anemia that are refractory to treatment with Erythropoietin (Epo). Recent work shows that early erythroid BFU-e and CFU-e progenitors form a developmental continuum characterized by gradual increase in the proportion of cells in S phase of the cycle. Here we proposed two distinct hypotheses to explain this finding, either the presence of quiescent progenitors or the gradual shortening of G1 and the cycle with differentiation. Using a mouse expressing a timer -protein transgene that reports cell cycle duration, we determined that, in vivo, early erythroid progenitors undergo orderly gradual shortening of the cycle as they mature and approach terminal differentiation. There was no evidence of quiescent BFU-e or CFU-e progenitors in tissue. We found that BFU-e and CFU-e progenitors are highly responsive to hypoxic stress and to its Epo and glucocorticoid mediators. Epo and hypoxia accelerated the pattern of gradual cell cycle shortening throughout early erythropoiesis, while conversely, dexamethasone prolonged the cycle specifically in proerythroblasts. Further, Epo and hypoxia generated rapid increase in early progenitor cell size and dynamic changes in cell surface marker expression. Our data suggest that high Epo or hypoxic stress promote rapid increase in the rate of growth in biomass across the entire erythroid trajectory including in the earliest BFU-e progenitors, and indicates that stress progenitors are of the same type and lineage as those sustaining basal erythropoiesis.

**Key Points:**

- A maturational process of gradual cell cycle shortening and increasing cell size in BFU-e and CFU-e is accelerated by Epo and hypoxia
- There are no quiescent BFU-e and CFU-e in tissue. Stress CFU-e arise from the same cell type and lineage as CFU-e in the basal state.

## Introduction

Erythropoiesis may be divided into an early erythropoiesis phase, and erythroid terminal differentiation (ETD). In early erythropoiesis, BFU-e and CFU-e progenitors undergo amplification and differentiation. A transcriptional switch transitions late CFU-e into ETD^1,2^, where erythroblasts (also known as erythroid precursors) undergo 3 to 5 maturational cell divisions before enucleating into reticulocytes.

Erythropoietin (Epo) and its receptor, EpoR, are essential regulators of erythropoiesis. Epo is used in the treatment of some types of anemia^3,4^ but is relatively ineffective in others, including myelodysplastic syndrome^5,6^ and Diamond Blackfan Anemia^7,8^. The pathology in the latter syndromes affects progenitors during early erythropoiesis, where the role of EpoR signaling is not well understood. Epo’s known cellular targets are late CFU-e and proerythroblasts, which depend on EpoR signaling for survival (**Fig. 1A**^9–11^). EpoR’s anti-apoptotic signaling in these cells is an established mechanism by which it regulates erythropoietic rate^12–16^. EpoR also regulates cycling in early ETD^17^. By contrast, Epo is dispensable for survival of BFU-e and early CFU-e^18^ and it is unclear whether and how it might regulate these progenitors. Erythroid progenitors integrate multiple extracellular inputs including SCF, glucocorticoids, IL-3, IL-6 and IL-17^19–27^. Of these, SCF, glucocorticoids and IL-17 either contribute to, or are essential for, increased erythropoietic rate in response to hypoxic stress^20,21,28^.

**Figure 1:**
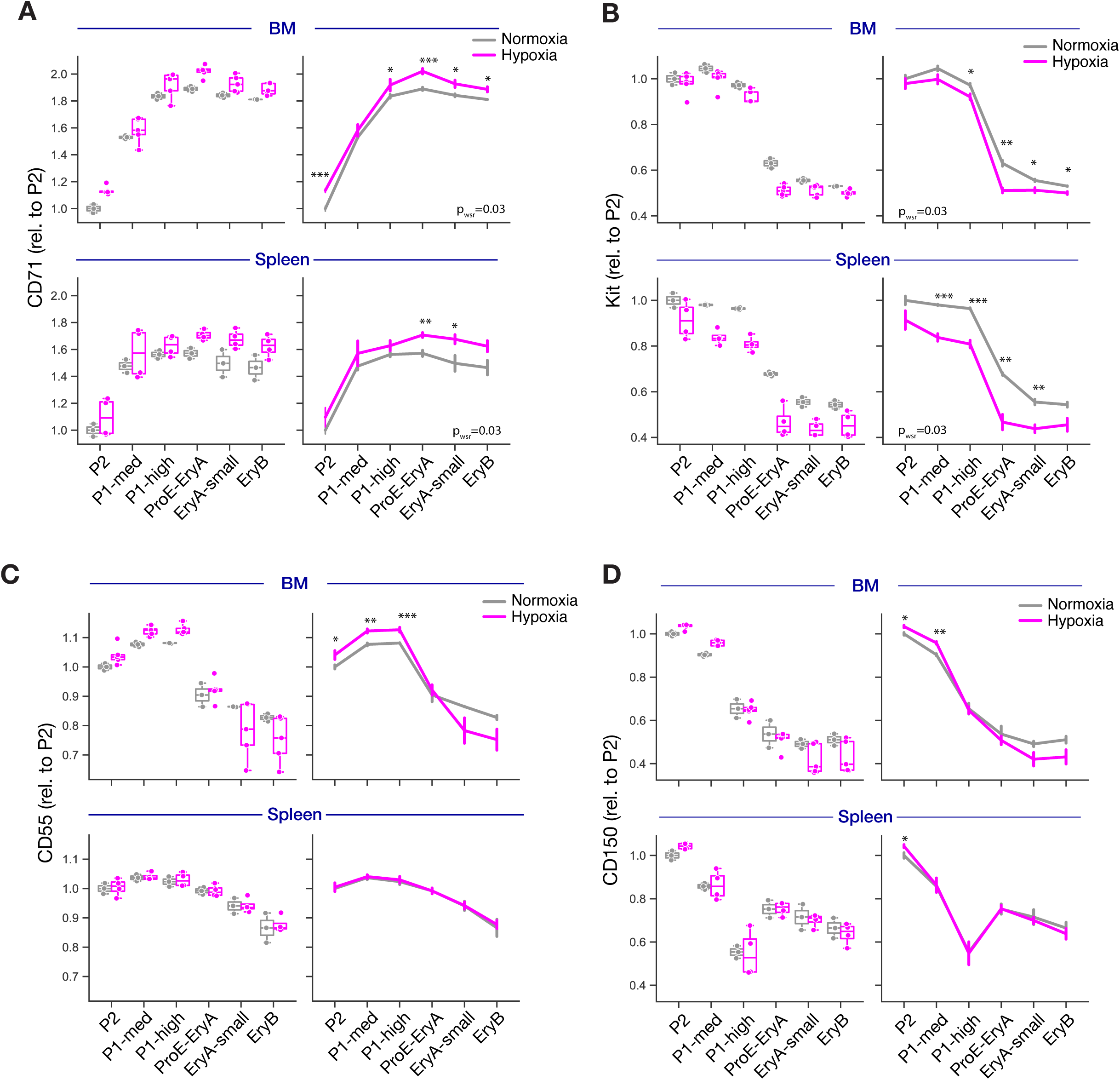
Hypotheses, predictions and experimental models. **A** Two alternative models to explain the observation that the proportion of S phase cells increases with early erythroid maturation. In Model 1, G1 shortening and hence cell cycle shortening accounts for the increase in the proportion of S phase cells. In Model 2, it is explained by a quiescent G0 subpopulation whose proportion decreases with maturation. **B** The H2B-FT transgenic mouse model ^37^. H2B-FT is under control of the rtTA doxycycline-inducible factor. H2B-FT fluoresces blue when first synthesized but matures into a long-lived red fluorescent protein within 1.2 hours. Blue fluorescence is unaffected by cell cycle duration, but red fluorescence increases over the length of the cycle. The ratio of Red to total fluorescence is a normalized measure of cell cycle duration. **C** Expected results for Model 1 and Model 2. In Model 1 (upper panel), all progenitors are in cycle, and form a monophasic distribution of cell cycle durations whose median decreases with maturation. In Model 2 (lower panel), the distribution of cell cycle durations at each maturation stage is biphasic, consisting of cycling cells whose cell cycle duration is unaffected by maturation, and of a quiescent population with predicted high levels of red /total fluorescence ratio whose proportion decreases with maturation. Violin lines denote median ± 25th and 75th percentiles. **D** Experimental design. Expression of H2B-FT is induced by adding doxycycline to the drinking water 2 weeks prior to the start of experiments. Mice were either placed in 10% oxygen or in normoxia and tissue was harvested at 24 hours for analysis. **E** Correspondence between the 10-color stain flow-cytometric gates and functional progenitor assignment ^1,29^. These FACS gates are used in all subsequent figures. **F, G** Representative flow cytometric analysis of tissue from normoxia and hypoxia using the 10-color stain. Panel F shows erythroblast gates corresponding to ETD ^14,15^. Panel G shows early erythroid progenitor gating strategy. The key to the functional progenitor subset for each gate ins in panel E.

The dearth of knowledge on the action of Epo in erythroid progenitors is in part due to the lack of a flow -cytometric strategy that identified these cells in tissue with high purity. In our recent work, we built on single-cell transcriptomics^1^ to develop a flow-cytometric strategy that identifies and stages all erythroid progenitors in mouse with high purity^1,29^, allowing analysis of their dynamic responses in vivo^1,20,30^.

Single-cell transcriptomics and flow-cytometric approaches show that erythroid progenitors follow a continuous process of differentiation. Starting at the multi-potential progenitor (MPP) stage, they acquire BFU-e and then CFU-e colony potentials, before activating ETD^1,27^. Although the maturational processes in early erythropoiesis are not well understood, it has become clear that they are intimately linked to cell cycle mechanisms. Single cell transcriptomics shows that early erythroid progenitor maturation is tightly correlated with a gradual increase in expression of S phase genes, including cyclin E, MCM helicase subunits and E2F4, all of which peak at the switch to ETD^1^. Functional analysis using a brief pulse of bromodeoxyuridine (BrdU) in vivo shows that the proportion of cells in S phase increases gradually during early erythroid progenitor maturation, from levels of ∼10% in early BFU-e, to ∼70-90% in late CFU-e/Proerythroblasts, accounting for the progressive increase in S phase genes ^1^. This striking increase in the proportion of S phase cells with differentiation stage was already deduced in early studies, based on increasing sensitivity to tritiated thymidine^31–33^. The underlying process that results in this pattern remains unclear. Here we propose two alternative hypotheses, also proposed by the authors of earlier studies^33^, that could explain this pattern (**Fig. 1A**). In the first (Model 1), the gradual increase in the proportion of S phase cells is the result of progressive shortening in G1 phase of the cycle as cells differentiate. In the second hypothesis (Model 2), the duration of both S phase and G1 remain constant throughout early erythropoiesis, but there is a subset of non-cycling, quiescent progenitors, whose proportion decreases as progenitors progress in their differentiation. A quiescent progenitor population might provide progenitor reserve that can be rapidly recruited during stress.

Indeed, recent reports of flow-cytometrically identifiable ‘stress CFU-e’ show that they preferentially expand in response to erythroid stress^30,34^. The cellular origins of stress progenitors is unclear, however. They may arise from the same lineage and cell type as steady-state progenitors, altering their cellular properties in response to extracellular stress factors; alternatively, as has been suggested for stress BFU-e^35^, they may arise from a distinct differentiation trajectory, with pre-programmed stress responses.

Here we used the novel progenitor flow-cytometric strategy (’10 color stain’^1,36^) to analyze erythroid progenitors in vivo in a recently-developed mouse model that is transgenic for a live-cell reporter of cell cycle duration^37^. This approach allowed us to determine why the proportion of S phase cells increases with progenitor maturation. We also investigated the effect of acute hypoxic stress, Epo and glucocorticoids on the abundance of erythroid progenitors in vivo, and on their cell cycle duration and cell surface marker expression. Our analysis suggests that cell cycle structure and speed is an intrinsic property dictated by erythroid maturational stage, as are cell size and the rate of growth in biomass. Further, extracellular stress factors like Epo and glucocorticoids have quite different effects on these progenitor parameters, suggesting distinct roles during stress, and potentially explaining their differing therapeutic properties in anemia.

## Methods

### Mice

Mice transgenic for H2B-FT (B6;129-*Gt(ROSA)26Sor^tm1(rtTA*M2)Jae^ Hprt1*^*tm2(tetO–mediumFT*)Sguo*^/Mmjax) were obtain from the Gao laboratory, Yale school of medicine. Eight to twelve week old male and female littermates were randomly assigned to treatment groups. This work was approved by the UMASS Chan Institutional Animal Care and Use Committee (IACUC) protocol PROTO202200017.

### Epo and dexamethasone treatment in vivo

The timer protein was induced by adding doxycycline to the drinking water 2 weeks prior to the start of experiments (1 g/liter). Mice were injected subcutaneously in a total volume of 4 microliters/g every 12 hours, with either Epo (0.25 IU/g , Procrit, Amgen), dexamethasone (2.5 mg/kg), both Epo and dexamethasone, or phosphate-buffered saline (PBS).

### Hypoxia treatment

Mice were placed for up to 48 hours in a BioSpherix A chamber (BioSpherix). Hypoxia (10% oxygen) was achieved by displacing oxygen with nitrogen at normal atmospheric pressure.

### Flow cytometry

Harvested spleen, bone marrow and blood were labeled with the indicated antibodies and analyzed on a spectral flow cytometer (Aurora, Cytek).

### Data analysis

Flow cytometry data was analyzed using FlowJo and the python programming language with matplotlib, pandas, seaborn, numpy and scipy packages.

**Additional methods** are provided in a supplemental file.

## Results

### Two hypotheses of early erythroid cell cycle behavior

The fraction of cells in S phase increases progressively during erythroid progenitor differentiation, while S phase duration remains largely constant^1^. We proposed two hypotheses to account for this behavior. In Model 1, erythroid progenitors undergo gradual shortening of both the G1 phase and the cell cycle, with S phase duration remaining constant. In Model 2, cell cycle duration remains constant, but not all progenitors are cycling; the fraction of quiescent progenitors decreases with differentiation (**Fig. 1A**). To distinguish these two models we used the ‘H2B-FT’ transgenic mouse ^37^, expressing histone H2B fused to a fluorescent timer (FT) protein under control of the doxycycline-inducible rtTA (**Fig. 1B**^37^). The FT protein domain of H2B-FT emits blue fluorescence when first synthesized, but matures over 1.2 hours into a long-lived red fluorescent protein. The ratio of blue to red fluorescence of H2B-FT was shown to report cell cycle duration in diverse cell types and tissues^37^. The ratio of blue protein to total (blue and red) fluorescence is a normalized index of cell cycle duration independent of protein synthesis rate (**Fig. 1B**^37^). We previously found that this reporter reproduces the known cell cycle shortening in ETD in vivo, confirming its utility^17^.

We grouped the continuum of early erythroid progenitors into three conventionally-recognized classes: BFU-e, early CFU-e and late CFU-e, based on flow-cytometric, transcriptomic and functional criteria^1^. For model 1, we expected to see a monophasic distribution of cell cycle durations in each of the progenitor classes, since they are all in cycle; the median duration for each of these distributions was expected to decrease with progression from BFU-e to early and then late CFU-e (**Fig. 1C**, top panel). For model 2, we expected a biphasic distribution for each progenitor class, since they each consist of both cycling and quiescent cells; the number of quiescent cells was expected to become proportionally smaller with each differentiation stage (**Fig. 1C**, lower panel).

### Early erythroid progenitors gradually shorten their cell cycle as they differentiate

To determine which of the two models describes early erythroid progenitors, we examined the H2B-FT mice in either normoxic conditions, or when placed in hypoxia (10% oxygen) for 24 hours (**Fig. 1D-G**). We identified progenitors in vivo using the ‘10 color stain’ (**Fig. 1E-G**^1,29^) where erythroid progenitors are Lin^−^Kit^+^CD55^+^CD49F^med/low^, and are further sub-divided into BFU-e (gate ‘P2’, **Fig. 1E,G**), early and late CFU-e (gates ‘P1-med’ and ‘P1-high’), based on increasing CD71 concomitantly with decreasing CD150 (**Fig. 1G).** Of note, the P2 gate contains all of the BFU-e in tissue, but also contains ∼ 25% of day 3 CFU-e^1^; for simplicity, below we refer to P2 as BFU-e. Erythroblasts in ETD were identified as ProE (Ter119^med^CD71^med/high^), EryA (Ter119^high^CD71^high^FSC^high^), EryB (Ter119^high^CD71^high^FSC^low^) and EryC (Ter119^high^CD71^low^FSC^low^, **Fig. 1E-F**^14,38^). EryA were subdivided into EryA-large and EryA-small, based on cell size (FSC)^14^. Of note, our prior analysis showed that S phase of the cycle is synchronized with the switch to ETD^1,2^. The ensuing initial upregulation of Ter119, corresponding to the ProE gate, coincides with G1 of the subsequent cycle. Consequently, ProE are enriched for G1, and the ensuing EryA-large gate is conversely enriched in S phase cells^2^. To obtain an asynchronously cycling population where all cell cycle phases are represented appropriately, we analyzed the union of the ProE and EryA-large subsets (’ProE-EryA’).

Cell cycle duration analysis in bone marrow and spleen showed monophasic distributions for all erythroid subsets, with the exception of EryC (**Fig. 2A**, **Fig. S1A**). Further, median cell cycle duration decreased with differentiation, with the shortest cycle taking place at the switch to ETD, corresponding to either late CFU-e (’P1-high’) or ProE (**Fig. 2A**). Conversely, cell cycle duration increased progressively during ETD. The biphasic distribution of cell cycle durations in EryC represents a mixture of cycling and post-mitotic late erythroblasts (**Fig. 2A**, **Fig. S1A**). The pattern obtained for EryC confirms that the H2B-FT transgene can accurately report the presence of mixed cycling and quiescent cell populations. The absence of biphasic populations in earlier subsets therefore shows that there are no quiescent erythroid progenitors in either bone-marrow or spleen. Taken together, our results support Model 1 as accounting for the cell cycle behavior of early erythropoiesis.

**Figure 2:**
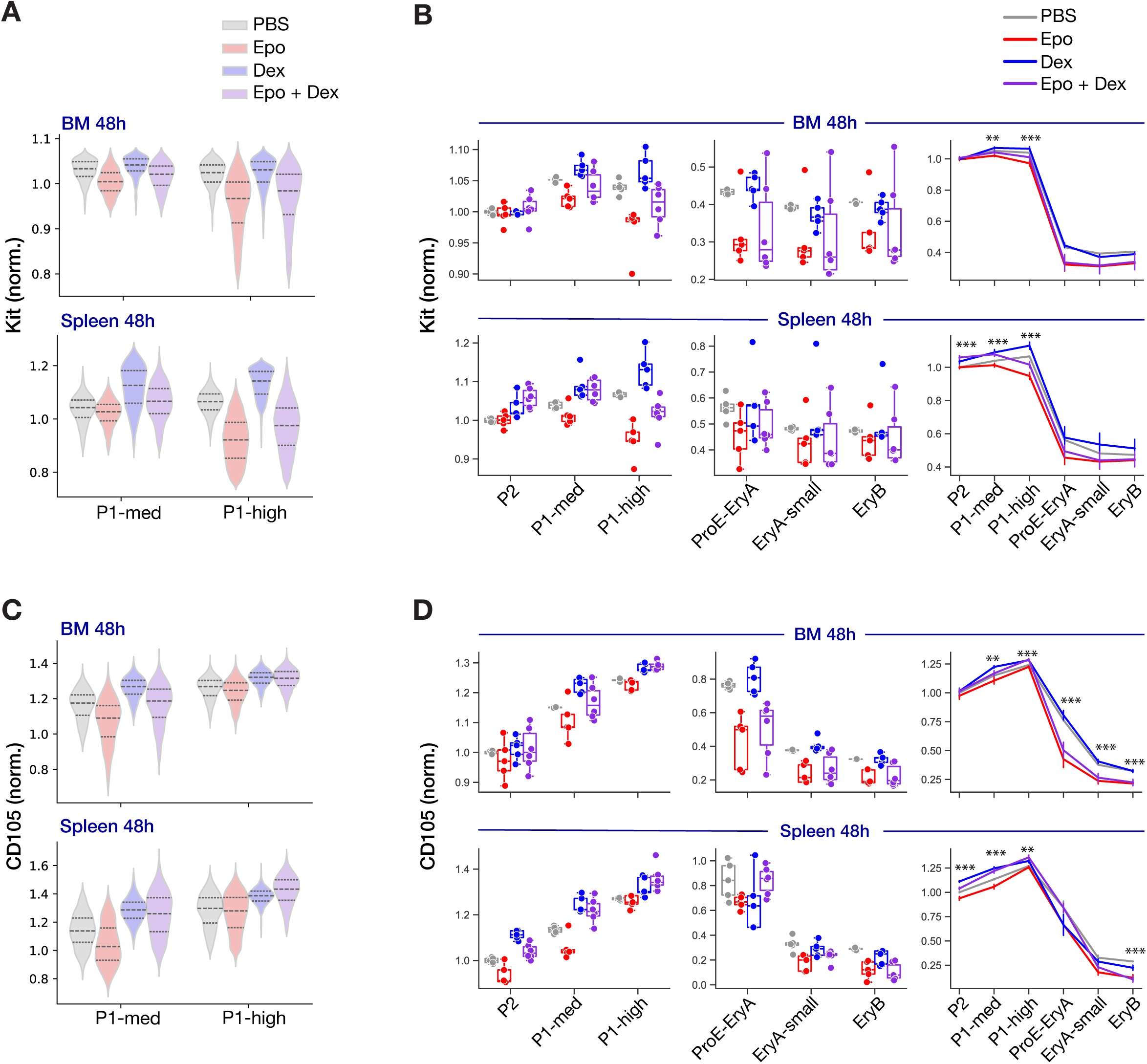
Gradual cell cycling shortening in maturing progenitors is accelerated by hypoxia. **A** Representative example of bone marrow and spleen early progenitor cell cycle durations. Violin plots are colored to correspond to progenitor maturational stage in Fig. 1A. Cell cycle length index was calculated as the ratio of red fluorescence to total blue and red fluorescence, and then expressed relative to this ratio in P2 (BFU-e). The y axis was cut off to allow visualization of spleen cycling progenitors but excludes some EryC cells that exited the cycle. The full dataset including all EryC cells is reproduced in **Fig. S1A**. Violin lines here and in all subsequent violin plots denote median ± 25th and 75th percentiles. **B** Distribution of cell cycle durations in the P1-high gate (late CFU-e subset) in individual mice. Each half violin corresponds to data from a single mouse in either normoxia or hypoxia. Pooled from 3 experiments. Cell cycle length was calculated as in panel A. **C** Comparison of cell cycle length distributions across all of the erythroid trajectory in spleen. Representative data from an experiment with two mice per conditions (either hypoxia or normoxia). **D** Cell cycle length across the entire erythroid trajectory for mice in either normoxia or hypoxia in bone marrow and spleen. Here and in all subsequent boxplots (left panels), data points correspond to the median value for a given progenitor subset in individual mice. Cell cycle length index was calculated as the ratio of red fluorescence to total blue and red fluorescence, and then expressed relative to the mean value for this ratio in P2 (BFU-e) in normoxia. Each box shows the median and central quartiles (25th to 75th percentiles), and whiskers extend by 1.5 times the inter-quartile range. Line plots (right panels) show the same dataset, with markers representing the mean ± sem value for all mice. Asterisks in grey (left panels) denote significant differences between the indicated subsets in normoxia (p-value for two-sided *t-test*; ***p<0.001, **p<0.01, *p<0.05). Asterisks in pink (right panels) denote significant differences between normoxia and hypoxia in individual subsets (with the same asterisk p-value notation). Right panels also show the p-value is for a Wilcoxon Signed Rank test (p_wsr_), a non-parametric paired test between normoxia and hypoxia across all differentiation stages in a given tissue. See all statistical analysis for this dataset in **Tables S1 and S2**.

### Hypoxic stress accelerates a pattern of gradual cell cycle shortening with differentiation

When mice were placed in 10% oxygen for 24 hours (**Figures 1D, F-G, 2B-C, S1B-D)**, the overall pattern of gradual cell-cycle shortening with erythroid differentiation was preserved, but at every differentiation stage, cell cycle duration in hypoxia was shorter than in normoxia (**Fig. 2B-D**). Hypoxia-induced cell cycle shortening was especially striking in spleen, including in the earliest, BFU-e progenitors (P2 subset, **Fig. 2D**). At each differentiation stage, the distribution of cell cycle durations shifted towards shorter cycles while remaining monophasic (**Fig. 2B-C**). This finding suggests a similar cell cycle response to stress by the entire erythroid progenitor population, and is against the presence of a sub-population of progenitors with unique stress-specific responses.

### Altered cell-surface marker expression in hypoxia suggests accelerated progenitor maturation

In hypoxia, progenitors expressed higher levels of CD71 and lower levels of Kit, changes that persisted through ETD (**Fig. 3A, B**). Both these trends are known consequence of enhanced EpoR signaling in erythroblasts and late CFU-e^39–41^ but were previously unknown in BFU-e and early CFU-e. Bone-marrow, but not spleen progenitors expressed significantly higher levels of CD55, a marker of erythro-megakaryocytic cell fate bias in MPPs (**Fig. 3C**^42^). CD150 levels increased in both spleen and bone-marrow BFU-e (**Fig. 3D**). We also noted increased cell size in bone-marrow BFU-e (**Fig. S1B**) and no significant changes in Ter119 or CD105 (**Fig. S1C-D**).

**Figure 3:**
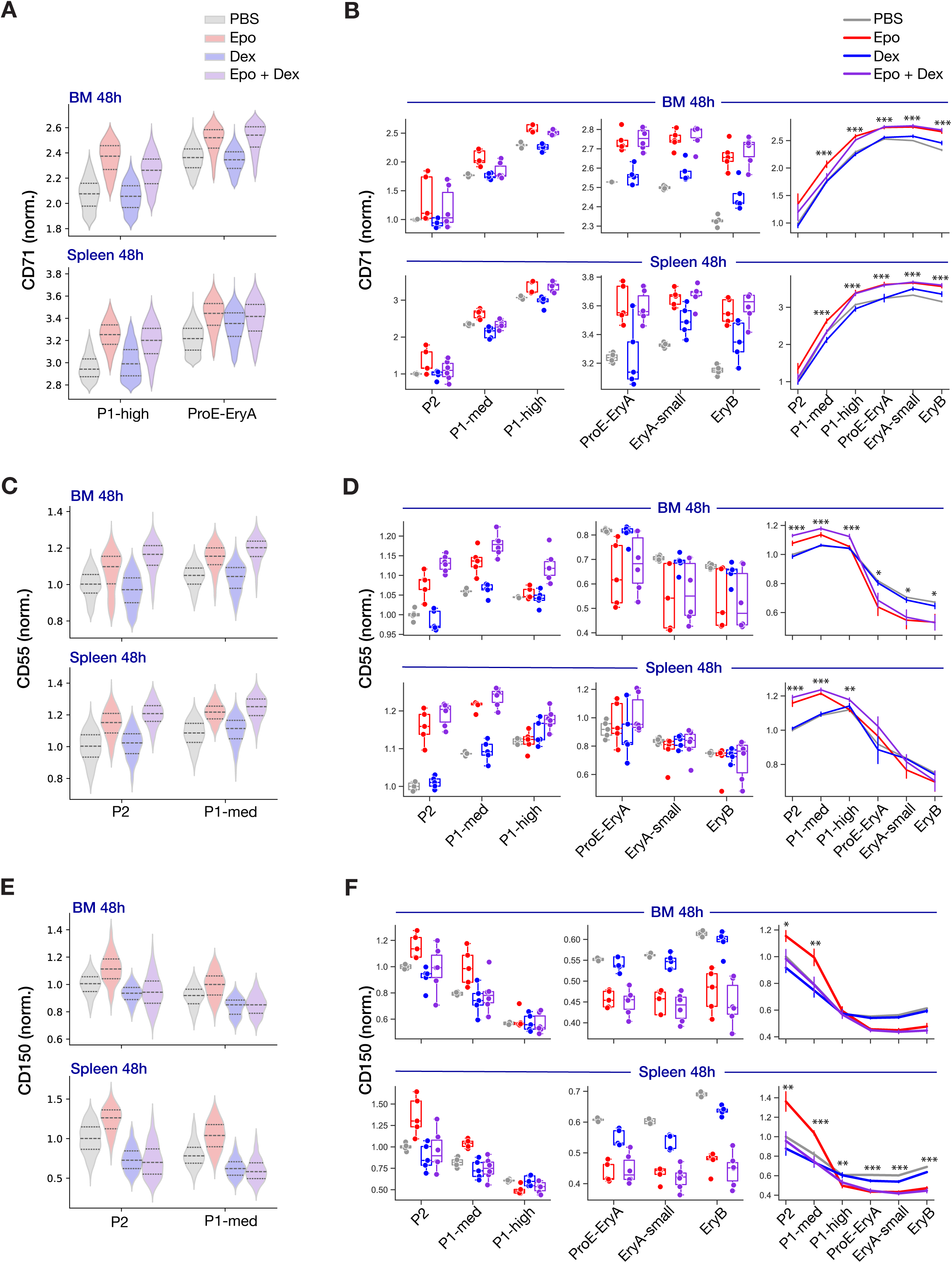
Cell surface marker dynamics in normoxia and hypoxia across the erythroid trajectory. **A-D** Levels of the indicated cell surface marker in each progenitor and erythroblast subset across the erythroid trajectory. Each cell surface marker is measured in arbitrary fluorescence units and expressed as a ratio, relative to the mean value for P2 in normoxia in the same experiment. Boxplots and line plots are as in the legend to Fig. 2D. Asterisks in grey (right panels) denote significant differences between normoxia and hypoxia in individual subsets (p-value for two-sided *t-test*; ***p<0.001, **p<0.01, *p<0.05). Right panels also show the p-value is for a Wilcoxon Signed Rank test (p_wsr_) between normoxia and hypoxia across all differentiation stages.

### No change in early progenitor number after 48 hours of Epo or Glucocorticoids

To investigate factors potentially responsible for progenitor cell cycle responses in hypoxia, we administered H2B-FT mice with two mediators of the erythropoietic stress response, Epo^15,16,43,44^ and glucocorticoids^19,28,45^. Mice were injected with either PBS, Epo (0.25 IU/g), the synthetic glucocorticoid dexamethasone (Dex, 2.5 mg/kg) or both Epo and Dex every 12 hours, and examined at 18 and 48 hours (**Fig. 4A**; n=4 and n= 6 mice per treatment at 18 h and 48 h respectively). There were no significant changes in either reticulocytes or spleen size at 18 hours. By 48 hours of Epo treatment, blood reticulocytes doubled (p= 0.026), as did spleen size (p=0.0002) (**Fig. 4B**), evidence of increased erythropoietic rate. Reticulocytes increased similarly in response to the combined Epo+Dex treatment (p=0.01), but spleen size did not increase; this reflects the shrinking effect that Dex has on the spleen, most likely the result of lymphocyte loss^46–48^. There was clear increase in spleen size when comparing Epo+Dex to Dex alone (p=10^−5^, **Fig. 4B**).

**Figure 4:**
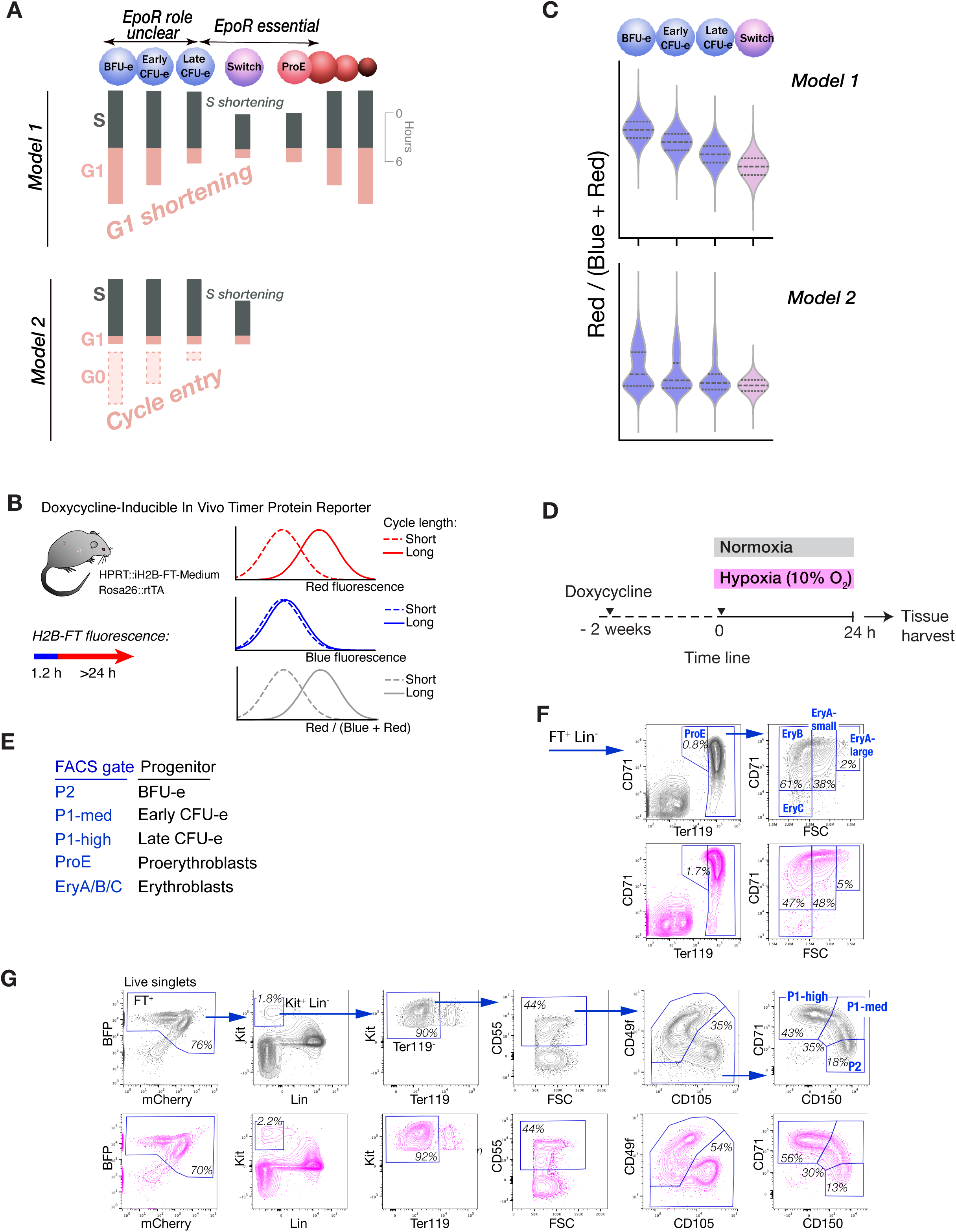
Effect of Epo and Dexamethasone administration on cell cycle length in vivo. **A** Experimental design. H2B-FT expression was induced by doxycycline 2 weeks prior to the start of experiments. Epo (0.25 IU/g), Dex (2.5 mg/kg), Epo+Dex or PBS were administered every 12 hours for up to 48 hours. Tissue was harvested and analyzed at 18 hours and 48 hours. **B** Spleen to body weight ratio (left panel) and reticulocytes in peripheral blood (right panel) at 48 hours. P-values are shown for one-way ANOVA for each panel, and in addition, for post-hoc pairwise analysis (independent two-tailed *t-test*). **C** Absolute cell number measured per g body weight for progenitors and precursors in each of the indicated subsets in bone marrow and spleen. Line plots are drawn to connect the mean ± sem of n=4 mice for each treatment at t=18 h and n=6 mice for each treatment at t=48 hours. Stars here and in all subsequent plots indicate the p-values (*p<0.05, **p<0.01, ***p<0.001) for one-way ANOVA between the 4 treatments at each progenitor stage, with false discovery rate (FDR) correction using the Benjamini-Hochberg (BH) procedure. P-values for post-hoc pairwise comparisons together with full descriptive and test data for all measured parameters are listed in **Tables S3 and S4**. **D** Cell cycle length distributions at each progenitor differentiation stage in spleen, for each of the indicated treatments. Representative data from a single experiment with 2 mice per treatment. **E** Summary data for cell cycle length for each of the four treatments from 3 pooled experiments. Datapoints, boxplots and line plots are as described in Fig. 2D. Grey asterisks in right panels denote significant differences in one-way-ANOVA for the given subset. Pairwise post-hoc t-test p-values are shown in left panels for Dex treatments in ProE-EryA (numerical values). Color-coded asterisks above box plots denote significant differences in post-hoc two-sided *t-test* for the indicated subsets for cytokine treatments vs. PBS (Epo: red, Dex: blue, Epo+Dex: purple). ***p<0.001, **p<0.01, *p<0.05. All other comparisons are in **Tables S3 and S4**. **F** Violin plots comparing cell cycle lengths in response to each of the four treatments in ProE-EryA, in each of 3 independent experiments, with 2 mice per treatment per experiment. Individual mouse data for the pooled experiments is shown in panel E.

By 48 hours, the number of late CFU-e and all ETD subsets in the spleen increased significantly in response to either Epo or the combined Epo+Dex treatment (**Fig. 4C**, p=0.0003, one-way ANOVA, see statistical tests for all individual pair-wise comparisons in **Tables S3, S4**). As example, late CFU-e and ProE-EryA increased 6 fold and > 10 fold respectively in response to either Epo or Epo+Dex; there was no increase in response to Dex alone. There was little change in the number of early progenitors (BFU-e and early CFU-e) in either tissue in response to Epo (**Fig. 4C**).

### Cell cycle duration modulation by Epo and Dex in erythroid progenitors

Although early progenitor number did not increase in 48 hours of Epo treatment, these cells may nevertheless have responded to Epo to support accelerated erythropoiesis, for example, by increasing flux through the early erythroid compartments. Indeed, we found significant shortening of cell cycle duration throughout the erythroid trajectory in response to either Epo or Epo+Dex, including significant shortening in BFU-e and early CFU-e in both spleen and bone marrow (**Fig. 4D-F, Tables S3, S4**). As in the response to hypoxia, the overall pattern of gradual cycle shortening with progenitor differentiation was preserved, and distributions of cell cycle duration for each progenitor stage remained monophasic (**Fig. 4D,E**).

Treatment with Dex alone had little effect on cell cycle duration, except in splenic ProE-EryA, corresponding to the time of the switch to ETD, where it prolonged the cycle (p=0.017 for Dex v. PBS, n= 6 mice, **Fig. 4E-F**). This finding is in agreement with the known effect of Dex on late CFU-e/Proerythroblasts in vitro, where it suppresses transition to ETD and promotes self-renewal cell divisions instead^45,49,50^, in part by preventing cell cycle shortening^51^. Of interest, the cell cycle response to combined treatment with Epo+ Dex resembles the response to Epo alone, suggesting Epo is the dominant regulator of the cycle with the treatment doses and timing we examined.

### Cell surface markers upregulated by Epo treatment: CD71, CD55 and CD150

Similar to the trend in hypoxia (**Fig. 3A**), treatment with either Epo or Epo+Dex resulted in the upregulation of CD71 significantly earlier in differentiation, starting in early CFU-e (P1-high, **Fig 5A, B**). There was also significant increase in the maximal levels of CD71 attained in ETD (**Fig 5A, B**). In the steady state, CD55 levels are high in BFU-e and increase further with differentiation, peaking in the P1 populations (CFU-e, **Fig. 3C**, **Fig. 5C,D**). Both Epo and Epo+Dex treatments induced earlier and higher levels of CD55, in BFU-e and early CFU-e (P2 and P1-med, **Fig. 5C,D**), resembling the effects of hypoxic stress (**Fig. 3C**). Finally, Epo but not the combined Epo+Dex treatment significantly increased CD150 in BFU-e and early CFU-e (p*_ttest_*=0.008 and 0.0001 for spleen BFU-e and early CFU-e, respectively, **Fig. 5E, F, Table S4**). Conversely, Dex as well as Epo+Dex decreased cell surface CD150 in the same progenitors, particularly in spleen. Indeed, this is one of the few examples in which it appears the response to the combined Epo+Dex treatment resembles the response to Dex, and there is a significant difference between Epo+Dex and Epo treatments (for Epo v Epo+Dex, p*_ttest_*=0.017 and 0.0009 for spleen BFU-e and early CFU-e, respectively, **Fig. 5E, F, Table S4**). We speculate on potential mechanisms of these cell surface marker changes in the Discussion section. Of note, cell surface marker distributions in each progenitor population remained monophasic in response to all treatments (**Fig. 5 A, C, E**).

**Figure 5:**
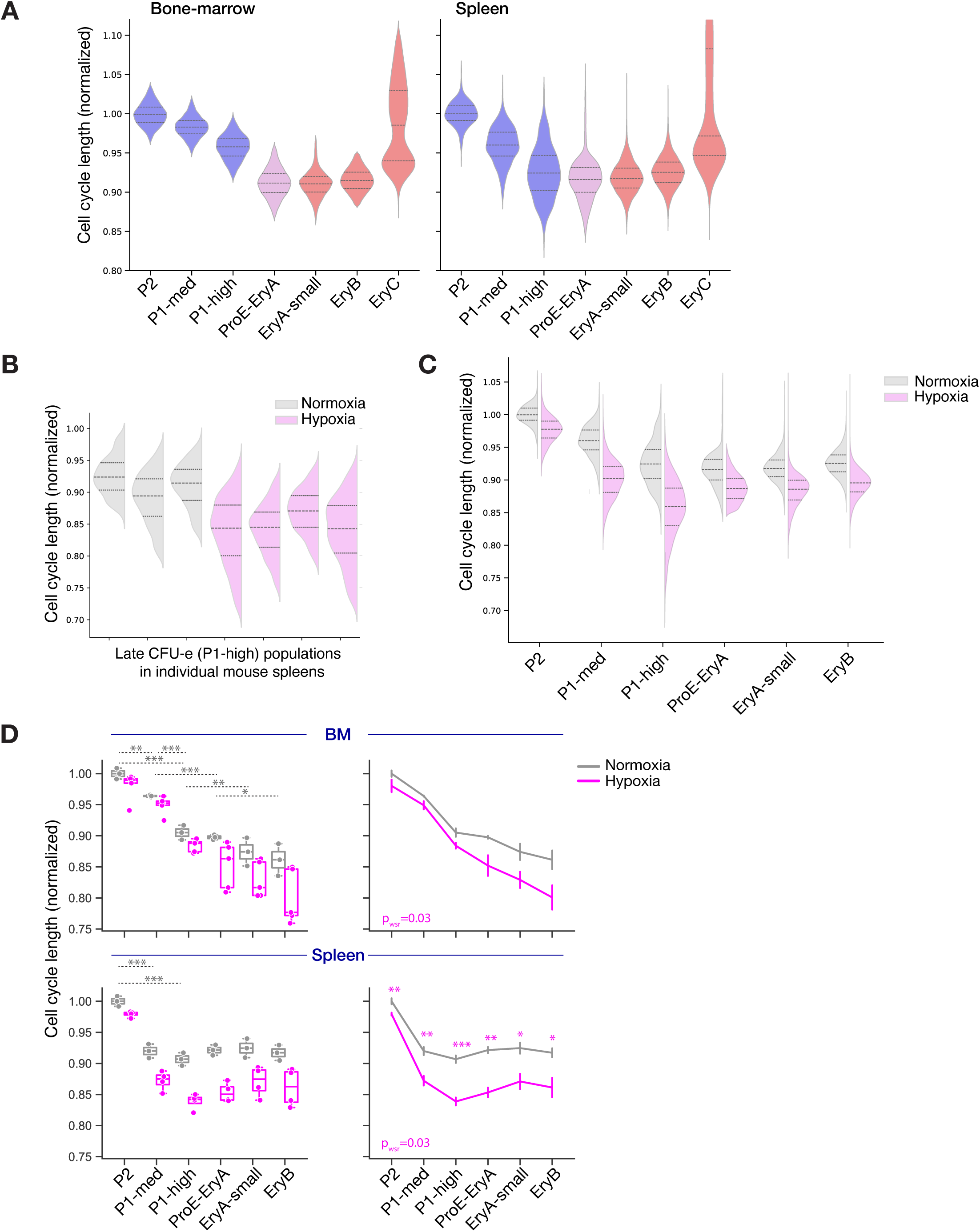
Rapid uprregulation of somes early progenitor cell surface marker in response to Epo. **A-F** Levels of the indicated cell surface marker in each progenitor and erythroblast subset across the erythroid trajectory. Each cell surface marker is measured as explained in the legend to Fig. 3. Boxplots and line plots are as in the legend to Fig. 2D. Panels A, C and E show violin plots for selected subsets in representative experiments, and panels B, D and F show summary data for the entire dataset. Statistical analysis and p-values as in the legend to Fig. 4C. Full statistical analysis in **Tables S3 and S4.**

### Opposing effects of Epo and Dex treatments: Kit and Endoglin expression

Dex and Epo altered Kit and Endoglin (CD105) expression in opposing ways, and the response to their combined action suggests that they are acting through independent pathways (**Fig. 6**). Kit increased in response to Dex, in all spleen progenitors and in bone marrow P1-med (**Fig. 6A, B, Table S4**). Conversely, there was significant decrease in Kit in response to Epo, in P1-high in spleen (p*_ttest_* = 0.00054) and in both P1 populations in the bone marrow (p*_ttest_* <0.01). The combined Epo+Dex treatment in the bone marrow nullified their individual effects. In the spleen, Dex predominated in earlier progenitors, increasing Kit (p<0.001 for P2, p< 0.02 for P1-med), whereas Epo dominated in late CFU-e, where Kit decreased. Endoglin was increased by Dex in all spleen progenitors, especially P2 (p*_ttest_* =6 × 10^−6^) and bone marrow P1 subsets (p*_ttest_* <0.001, **Fig. 6C, D, Table S4**). Epo decreased endoglin in multiple subsets, including spleen progenitors P2 P1-med (p*_ttest_*∼0.015) and both spleen and bone marrow ETD. In the response to the combined treatment, Dex predominated in progenitors, increasing endoglin in all spleen progenitors and in bone marrow P1-high (p*_ttest_* <6 × 10^−5^); Epo dominated in ETD, decreasing endoglin in both bone-marrow and spleen.

**Figure 6:**
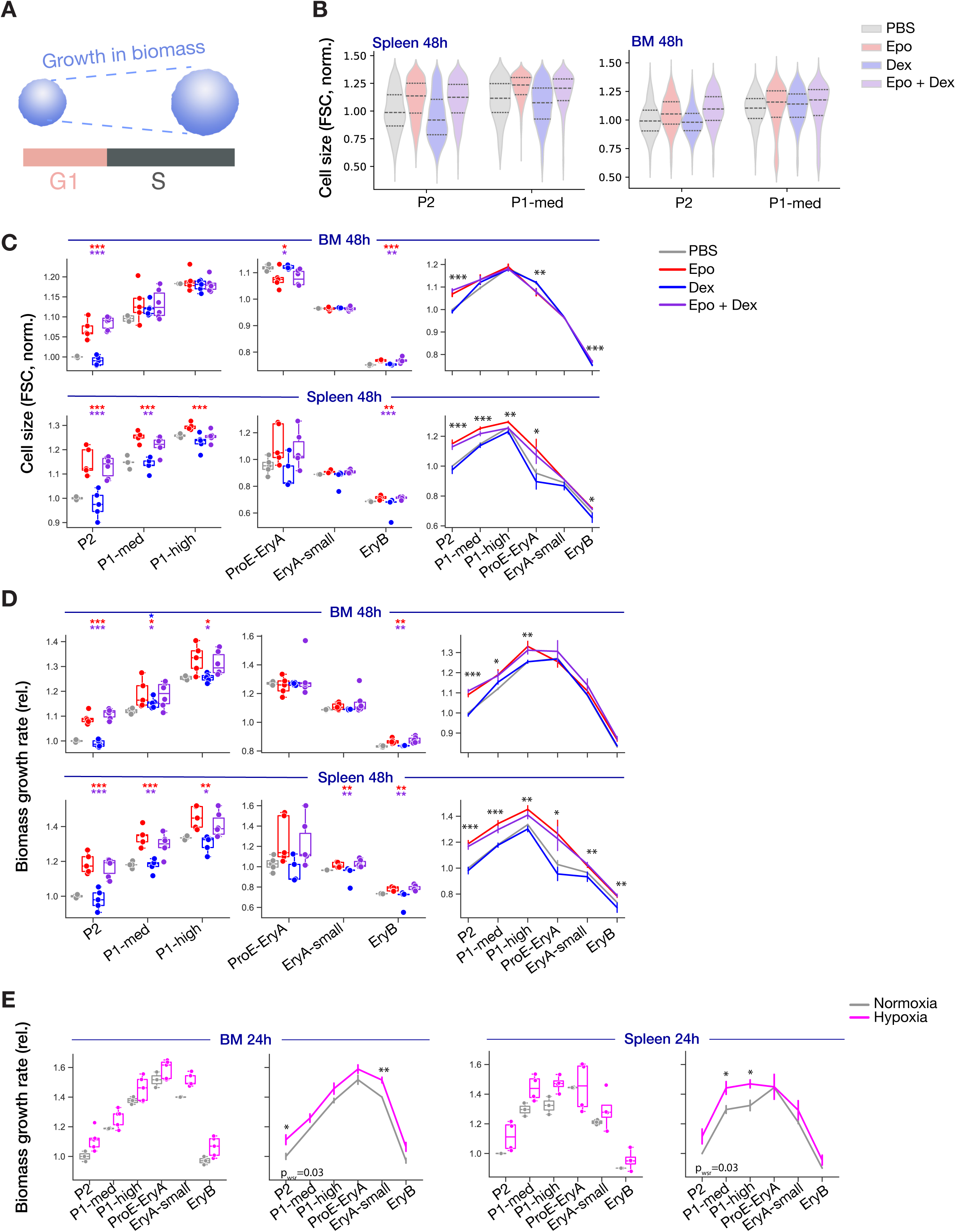
Change in Kit and Endoglin expression in early erythroid progenitors in response to Epo and Dex. **A-D** Levels of the indicated cell surface marker in each progenitor and erythroblast subset across the erythroid trajectory. Data presentation and statistics as described in the legend to Fig.5.

### Progenitor cell size and rate of growth in cell biomass are increased by Epo and hypoxia

To maintain cell size over cell generations, cells need to double their size during each cycle (**Fig. 7A**). Cell cycle shortening may be expected to result in smaller cell size, since less time is available for growth in cell biomass. Given the cell cycle shortening throughout erythropoiesis in response to Epo (**Fig. 4D-E**), we examined cell size using the flow cytometric forward scatter parameter (FSC). In the steady state, cell size increased gradually during early erythropoiesis, peaking in late CFU-e (P1-high, **Fig. 7B-C**). As is well established, cell size then declined throughout ETD (**Fig. 7C**). In response to Epo or Epo+Dex, there was significant increase in cell size, in spite of accelerated cell cycle shortening, particularly in BFU-e (bone marrow and spleen), early and late CFU-e (in spleen, **Fig. 7B-C**). In agreement with our previous work, Epo also increased cell size in ETD (**Fig. 7C**)^17^.

**Figure 7:**
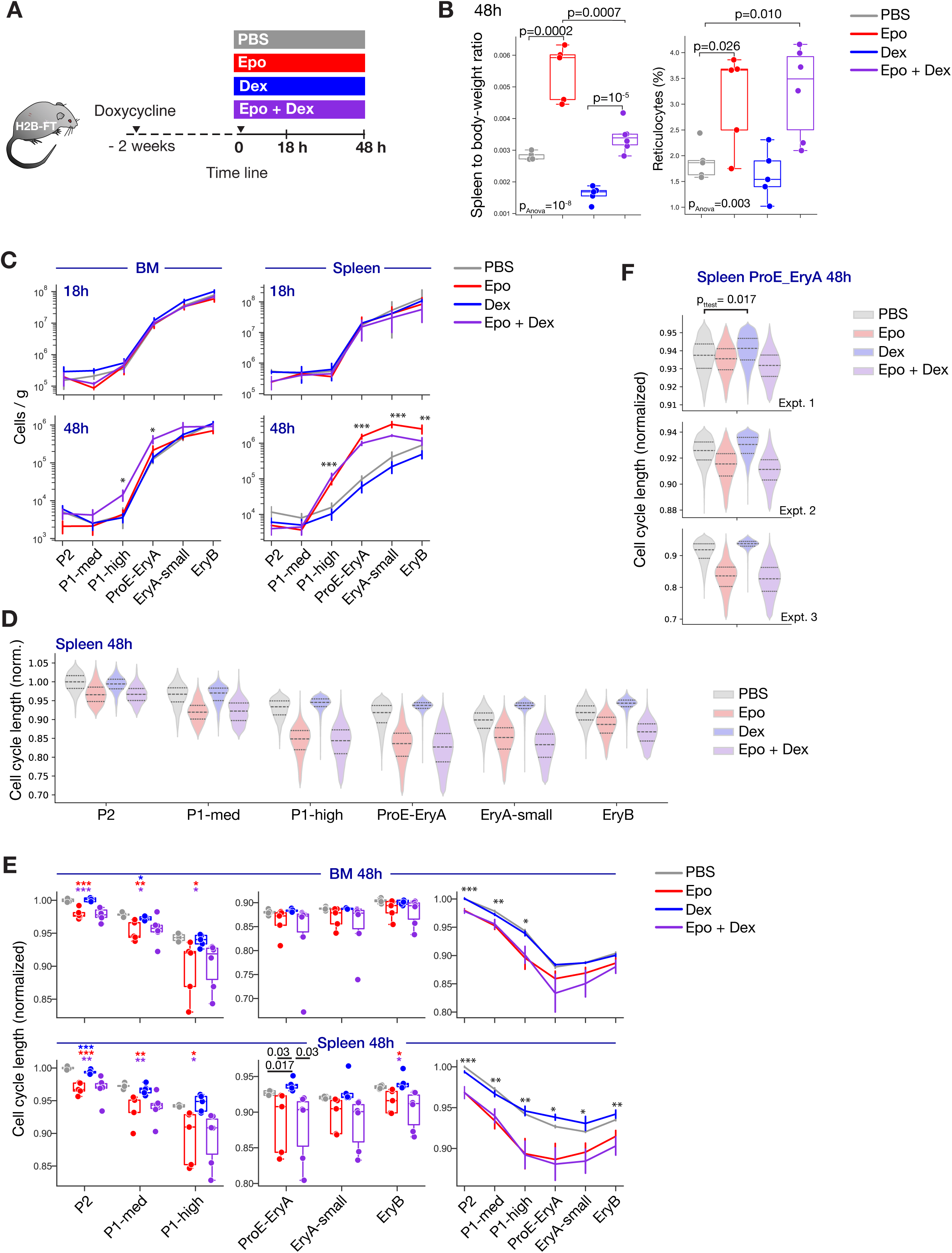
Epo accelerates the rate of growth in biomass throughout the erythroid trajectory. **A** Cartoon illustrating that cell size is a function of the rate of growth in biomass and the length of the cell cycle. **B** Cell size distributions in early progenitor subsets in a representative experiment, showing increased cell size in response to Epo or Epo+Dex. **C-E** Cell size and biomass growth rates following Epo, Dex or Epo+Dex treatment (C,D) or in hypoxia (E). Boxplots, line plots and p-values as in the legend to **Fig 2D**. Biomass growth rate index was calculated as the ratio of cell size (FSC units) to cell cycle length (fluorescence ratio). Asterisks above line-plots and boxplots indicate statistical significance as described in **Fig 4E**.

To determine what the simultaneous change in cell size and cell cycle duration imply regarding the rate of growth in biomass, we calculated a growth rate index, by dividing cell size (in FSC units) by cell cycle duration (in fluorescence ratio units) for each subset (**Fig. 7D**). This index suggested that the rate of growth in biomass increases throughout early erythropoiesis, peaking in late-CFU-e or ProE. Treatment with either Epo or Epo+Dex accelerated the rate of growth in biomass significantly, throughout erythropoiesis in the spleen, and in bone-marrow progenitors (**Fig. 7D**). Similarly, there was significant acceleration in cell biomass growth in hypoxic stress (**Fig. 7E**). Of note, we may be underestimating the increase biomass growth, since the changes in cell size and in cell cycle length through the H2B-FT reporter may be slow to reflect the underlying process.

## Discussion

Relatively little is known of the behavior of erythroid progenitors in vivo. The present work shows that progenitors undergo well-ordered, continuous phenotypic change: cell cycle shortening, increasing cell size and accelerating growth in biomass. These and a number of cell-surface markers undergo further rapid change in response to Epo and stress. The response of erythroid progenitors to dexamethasone differs in significant respects to their response to Epo, potentially explaining why Epo and glucocorticoids differ in their therapeutic effectiveness in distinct types of anemia. For all measured parameters, their distribution at each differentiation stage was monophasic and remained so in response to stress. These observations suggest that stress progenitors are of the same type and lineage as progenitors that support steady-state erythropoiesis.

Early erythropoiesis is marked by gradual increase in the proportion of S phase cells^1,31–33^. Here we examined two hypotheses that could explain these findings: either gradual shortening in G1 and the cell cycle, or the presence of quiescent early progenitor subpopulations that decrease with differentiation.

Unlike hematopoietic stem cells (HSCs) or MPPs, which were shown to include both cycling and quiescent populations^52^, our findings support the first hypothesis (Model 1, **Fig. 1A**) and suggest that there are no quiescent BFU-e or CFU-e in tissue. Given the monophasic distributions of CD71, Kit, CD55, CD150, CD105 and cell size in the steady state and in response to stress, we conclude that stress CFU-e, recently identified as CFU-e that undergo preferential expansion in response to stress^30,34^, originate from the same cell type and lineage as steady-state progenitors. All of the erythroid progenitors in tissue therefore appear to have the capacity to respond to stress, and the extent to which they do so is distributed monophasically at each differentiation stage. These findings are in agreement with single-cell transcriptomics, which lacks evidence for distinct sub-types of Epo-responsive progenitors^1^, and with a study in which stress-induced HbF-expressing erythroblasts could not be distinguished from “A” erythroblasts either transcriptionally or by proteomics^53^.

Epo -mediated cell cycle shortening is more pronounced in spleen than in bone-marrow progenitors, consistent with the spleen being an established site of erythropoietic expansion during stress in mice^31,32,54–56^. Spleen also differed from bone-marrow in expressing higher CD150 in BFU-e in response to Epo, more pronounced downregulation of Kit in response to either Epo or hypoxia, higher endoglin in response to Dex, and a more significant increase in cell size and rate of growth in biomass across many progenitor and precursor subsets in response to stress. Epo-mediated pro-survival functions, such as its suppression of Fas and FasL, are also more pronounced in the spleen^14,57^. The mechanism making spleen erythroid cells more responsive to Epo and stress is unclear. Cells undergoing erythropoietic expansion in the spleen are thought to originate in BFU-e that migrate from bone marrow with the onset of stress^31^. It is likely therefore that differences between spleen and bone-marrow progenitors’ responses are not cell autonomous, and may possibly reflect differences in the erythropoietic niche.

Treatment with Epo or Epo+Dex for 48 hours induced a robust expansion in late CFU-e and in ETD cells, but there was no increase in BFU-e or early CFU-e number. This may be due to the difficulty of detecting significant change in rare progenitor populations with high variance at an early stage of stress. Alternatively, increased flux through these compartments may take place without an increase in cell number. Our results are consistent with the latter possibility. We found that BFU-e and early CFU-e progenitors respond dynamically and rapidly to Epo and hypoxia, by altering cycle duration, cell size and cell surface marker expression. This includes increasing CD150 levels in response to Epo in BFU-e, but decreasing CD150 in response to Dex (whether alone or in combination with Epo). CD150 is an HSC marker. By analogy with label-retaining cell experiments, we speculate that it is lost passively as these cells divide and differentiate. Its higher levels following Epo treatment may indicate higher flux from HSCs and MPPs into the BFU-e compartment, and/or faster flux through the BFU-e compartment, with fewer cell divisions taking place and hence higher levels of CD150 retained. Conversely, glucocorticoids are established as slowing erythroid maturation^27,45^, possibly leading to more cell divisions at each differentiation stage and therefore to loss of CD150 at earlier stages of differentiation. Evidence that Epo accelerates maturation during early erythropoiesis also comes from the finding that at each differentiation stage, Epo enhanced CD71 upregulation, loss of Kit and shortening of the cycle, all changes that occur as progenitors differentiate. Acceleration of these processes by Epo might perhaps accelerate the maturational process. By contrast, Dex upregulated Kit, a receptor required for sustaining earlier progenitors. Dex also prolonged the splenic ProE cycle, a potential mechanism for slowing erythroid maturation and the onset of ETD^51^, and increased Endoglin expression in progenitors. Endoglin is a TGFβ co-receptor and its promoter contains a glucocorticoid response element (GRE)^58^, suggesting that it may be a target of glucocorticoid signaling in erythroid progenitors.

Of note, although Epo shortens the cycle in both early erythropoiesis and in ETD, the consequences for maturational speed and cell cycle number may differ during these two phases of erythropoiesis. In ETD, where maturation is associated with prolongation of the cycle (**Fig 1A**), we reported that Epo-induced cycle shortening is associated with more numerous cell divisions and prolonged maturation^17^.

Conversely, our work here suggests that in early erythropoiesis, where maturation is associated with cell cycle shortening, further cycle shortening by Epo appears to accelerate the maturational process, possibly also reducing the number of associated cell divisions. We speculate that Epo’s opposing effects on maturational speed and on cell cycle number in early erythropoiesis versus ETD may account for the observation that Epo increases the levels of CD150 and CD55 in early erythropoiesis, but decreases their levels in ETD (Figs 5D,5F). Thus, assuming these cell surface markers are relatively stable, their levels would be reduced by the dilutional effect of cell division. In early erythropoiesis, Epo induces faster but fewer cycles, allowing more of these markers to be retained on the cell surface. By contrast, in ETD, Epo induces faster and more numerous cycles, resulting in loss of these cell surface markers at an earlier maturational stage.

Both Epo and hypoxia decreased Kit on erythroid progenitors. This finding may be surprising, since Kit and EpoR signal cooperatively and synergize in the erythroid stress response^21,49,54,59,60^. Our results are consistent, however, with Epo decreasing Kit on leukemic erythroid cells^41^, and with Kit over-expression in progenitors when EpoR is deleted (MS, in preparation). Although EpoR and Kit both stimulate proliferation of late CFU-e/proerythroblasts, they have opposing effects on the rate of progenitor differentiation. Thus, constitutively active Kit inhibits progenitor terminal differentiation [12], whereas here we find that Epo may accelerate progenitor differentiation. These opposing effects may be viewed as alternative pathways for increasing erythropoietic rate. High erythropoietic rate requires both a large pool of progenitors, and their fast differentiation. These two requirements are in conflict: accelerated differentiation depletes the progenitor pool, and conversely, progenitor pool expansion is at the expense of fast differentiation. The optimal balance between these two effects presumably depends on whether the required increase in erythropoietic rate is acute or chronic. These dynamics are likely to be regulated by the balance between Epo, accelerating progenitor differentiation, and Kit and glucocorticoids, which slow differentiation down. Future work will be required to further elucidate these effects in erythroid progenitors during stress.

## Acknowledgements

The authors thank the staff of the UMASS Chan flow cytometry core for their help. This work was supported by the National Institutes of Health R01DK120639 and R01 DK136321 to MS and by NIH S10OD028576 to UMASS Chan Medical school, for a BDFACSFusion.

## Authorship Contributions

AW, LL and DN designed and performed experiments, AW, LL and MS analyzed the data and wrote the manuscript.

