## Supplemental Figures and Methods for "Epo and hypoxia accelerate a pattern of gradual cell cycle shortening in BFU-e and CFU-e erythroid progenitors *in vivo*"

short title: Cell cycle shortening in erythroid progenitors

Ashley Winward[1], Logan Lalonde[1], Divya Nair[1], Merav Socolovsky[1#]

Content:

1. Supplemental Figures

2. Supplemental Methods

### Supplemental Figures

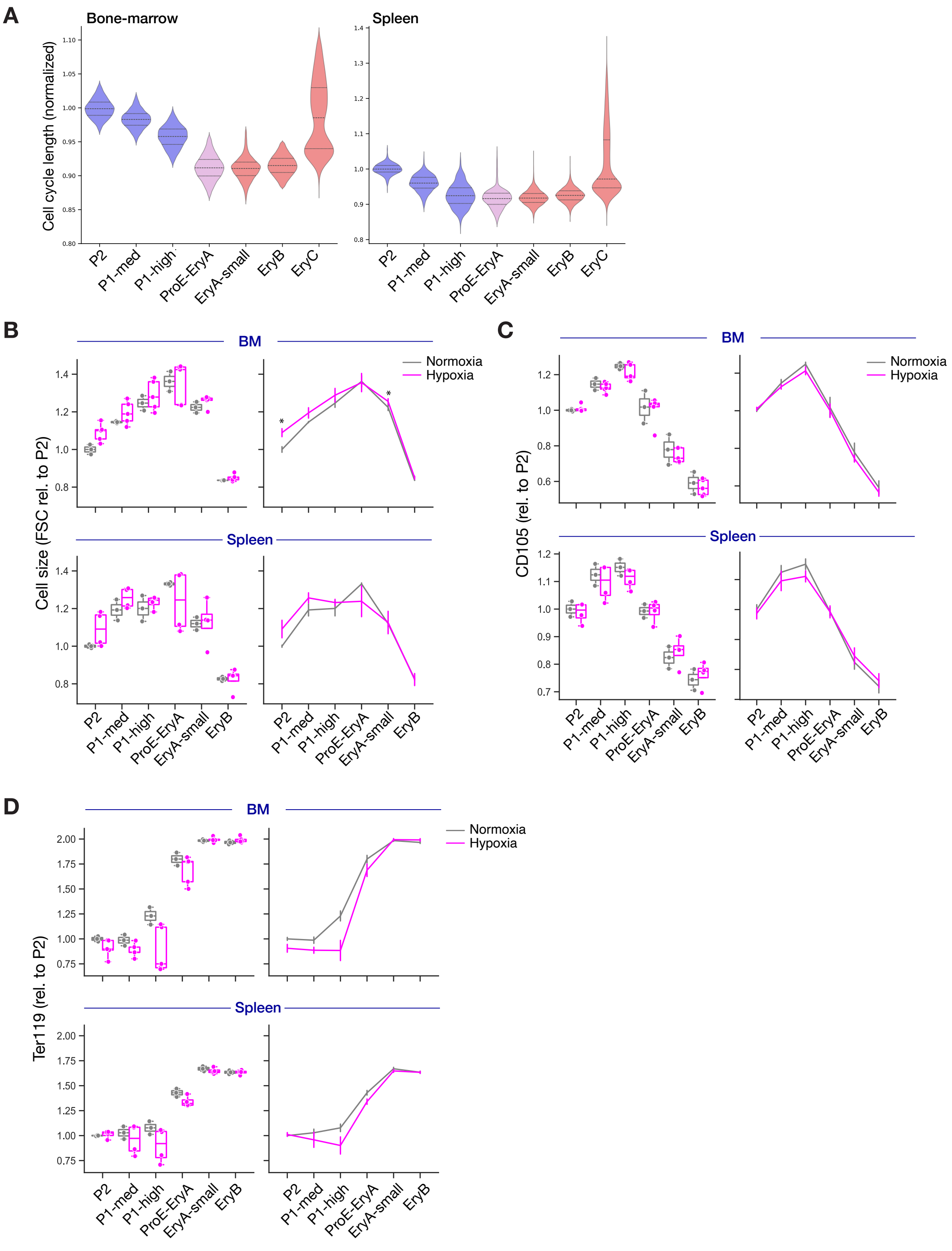


**Figure S1: Effect of hypoxia on cell cycle duration and on cell surface markers**

**A** Representative example of bone marrow and spleen early progenitor cell cycle durations. The same data as in **Fig 2A**, showing all of the EryC data in the spleen.

**B-D** Levels of the indicated cell surface markers in hypoxia or normoxia. Boxplots and line plots as described in **Fig. 2D**.

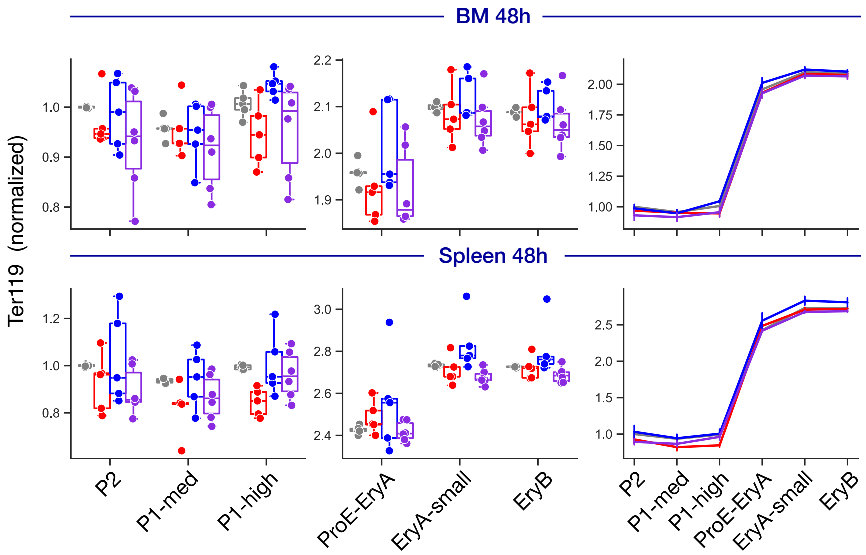


**Figure S2: No change in Ter119 expression with Epo or Dex treatments**
Ter119 in response to treatment with Epo, Dex, or Epo+Dex. Boxplots and line plots as described in **Fig. 2D**.

**Supplemental Methods**

**Antibody staining for flow cytometry**

Antibody labeling was done on freshly harvested blood, bone marrow or spleen cells at 4°C. All antibody panels included either Fc block or rabbit IgG (Jackson ImmunoResearch Laboratories, Item# 011-000-003). All cells were post-labeled with DAPI (Thermofisher Scientific, Item # D1306) or LIVE/DEAD Aqua (Invitrogen, Item # L34957) to exclude dead cells.

**Reticulocytes** were measured by flow-cytometric analysis of blood labeled with the DNA stain Draq5 (Cell Signaling Technology, Item # 4084L) and with CD71 antibody (Biolegend, Clone RI7217).

**Erythroid precursors** (ProE, EryA/B/C) were identified in spleen and bone marrow samples that were labeled with a lineage marker cocktail (Gr, AF700, Biolegend, RB6-8C5; Mac1, AF700, Biolegend, M1/70 Clone; CD4, AF700, Biolegend, RM4-5 Clone; CD8a, AF700, Biolegend 53-6.7 Clone; CD19, AF700, Biolegend, 1D3/CD19 Clone; F4/80, AF700, Biolegend, BM8 Clone) and with antiobodies to CD71 (PeCy7, Biolegend, RI217 Clone) and Ter119 (BUV395, BD Biosciences, Ter119 Clone).

**Early erythroid progenitors** (BFU-E/P2, early CFU-E/P1-medium, late CFU-E/P1-high) were identified by labeling with antibodies directed at Kit (APC-Cy7, Biolegend, 2B8 Clone), lineage marker cocktail as above, Ter119 (BUV395, BD Biosciences, Ter119 Clone), CD71(PeCy7, Biolegend, RI217 Clone), CD55 (AF647, Biolegend, RIKO-3 Clone), CD49f (AF488, Biolegend, GoH3 clone), CD105 (PE, Biolegend, MJ7/18 Clone), CD150 (BV650, Biolegend, TC15-12F12.2 Clone), CD41(BV605, Biolegend, MWReg30 Clone) (the '10 color panel').

**Single color controls** for mCherry and blue fluorescent protein (BFP) were prepared as follows : antibodies for mCherry (Thermofisher, 16D7 Clone) or BFP (Isbio, 3A6 Clone) were incubaed with UltraComp eBeads™ (Invitrogen item #01-2222-42). Antibody bound beads were then incubated with recombinant BFP (abcam item# ab285700) or mCherry (abcam item# ab199750).
